# Reference genome and demographic history of the most endangered marine mammal, the vaquita

**DOI:** 10.1101/2020.05.27.098582

**Authors:** Phillip A. Morin, Frederick I. Archer, Catherine D. Avila, Jennifer R. Balacco, Yury V. Bukhman, William Chow, Olivier Fedrigo, Giulio Formenti, Julie A. Fronczek, Arkarachai Fungtammasan, Frances M.D. Gulland, Bettina Haase, Mads Peter Heide-Jorgensen, Marlys L. Houck, Kerstin Howe, Ann C. Misuraca, Jacquelyn Mountcastle, Whitney Musser, Sadye Paez, Sarah Pelan, Adam Phillippy, Arang Rhie, Jacqueline Robinson, Lorenzo Rojas-Bracho, Teri K. Rowles, Oliver A. Ryder, Cynthia R. Smith, Sacha Stevenson, Barbara L. Taylor, Jonas Teilmann, James Torrance, Randall S. Wells, Andrew Westgate, Erich D. Jarvis

## Abstract

The vaquita is the most critically endangered marine mammal, with fewer than 19 remaining in the wild. First described in 1958, the vaquita has been in rapid decline resulting from inadvertent deaths due to the increasing use of large-mesh gillnets for more than 20 years. To understand the evolutionary and demographic history of the vaquita, we used combined long-read sequencing and long-range scaffolding methods with long- and short-read RNA sequencing to generate a near error-free annotated reference genome assembly from cell lines derived from a female individual. The genome assembly consists of 99.92% of the assembled sequence contained in 21 nearly gapless chromosome-length autosome scaffolds and the X-chromosome scaffold, with a scaffold N50 of 115 Mb. Genome-wide heterozygosity is the lowest (0.01%) of any mammalian species analyzed to date, but heterozygosity is evenly distributed across the chromosomes, consistent with long-term small population size at genetic equilibrium, rather than low diversity resulting from a recent population bottleneck or inbreeding. Historical demography of the vaquita indicates long-term population stability at less than 5000 (*Ne*) for over 200,000 years. Together, these analyses indicate that the vaquita genome has had ample opportunity to purge highly deleterious alleles and potentially maintain diversity necessary for population health.

## Introduction

In the afternoon of November 4, 2017, an adult female vaquita porpoise (*Phocoena sinus*), the smallest and rarest cetacean in the world, was captured in a massive effort to save the species by bringing into captivity as many as possible of the estimated maximum of 30 remaining individuals at the time (Thomas et al., 2017). This represented only the second live capture of a vaquita ever, the first of which, just a few weeks earlier, resulted in release of the animal after only hours when it showed signs of continuing stress. Despite the efforts of an international team of scientists and experts in porpoise capture and care, the second captured vaquita (V02F), suffered stress-induced cardiac failure and died approximately seven hours after initial capture (Rojas-Bracho et al., 2019). That death ended the effort by the Vaquita Conservation, Protection, and Recovery (VaquitaCPR) project to temporarily protect vaquita near their native habitat in the northern Gulf of California, near San Felipe, Mexico. However, the careful planning and presence of veterinarian experts in marine mammal stranding response allowed for an immediate necropsy that went through the night, with harvest and storage of ovaries and other tissues for delivery to facilities 260 miles north near San Diego, California for tissue culture and cryopreservation. By eight p.m. the next day, within 24 hours of the animal’s cardiac arrest, the tissues were delivered to the Institute for Conservation Research, San Diego Zoo Global, for the culture of cells from as many tissues as possible. After weeks of tissue culture, cells were harvested and banked for future research, and frozen samples sent to the Vertebrate Genome Lab at The Rockefeller University to extract ultra-high molecular weight DNA and RNA for genome sequencing, assembly and transcriptome annotation.

This extraordinary effort to extract as much information as possible from the VaquitaCPR project reflects the broad scientific value placed on biodiversity and conservation. Sequencing of reference genomes is increasingly recognized as an important contribution to identify, characterize and conserve biodiversity (Garner et al., 2016; Harrisson, Pavlova, Telonis-Scott, & Sunnucks, 2014; He, Johansson, & Heath, 2016; Kraus et al., submitted; Morin et al., in revision; Supple & Shapiro, 2018), especially for species that are naturally rare and difficult to study. Reference genomes provide primary data to understand evolutionary relationships (Arnason, Lammers, Kumar, Nilsson, & Janke, 2018; Zhou et al., 2018), historical demography (Armstrong et al., 2019; Andrew D Foote et al., 2016; Morin et al., 2018a; Robinson et al., 2016; Westbury, Petersen, Garde, Heide-Jorgensen, & Lorenzen, 2019), evolution of genes and traits (Autenrieth et al., 2018; Fan et al., 2019; A. D. Foote et al., 2015; Morin et al., in revision; Springer et al., 2016a; Springer, Starrett, Morin, Hayashi, & Gatesy, 2016b; Yim et al., 2014) and susceptibility to inbreeding and outbreeding depression (Chattopadhyay et al., 2019; Hedrick, Robinson, Peterson, & Vucetich, 2019; Robinson, Brown, Kim, Lohmueller, & Wayne, 2018; Tunstall et al., 2018). Genomic resources also provide the tools for broader studies of population structure, relatedness and potential for recovery (e.g., Garner et al., 2016; Morin et al., 2018b; Tunstall et al., 2018).

The vaquita was described for the first time in 1958 (Norris & McFarland, 1958) and has been characterized as a naturally rare endemic species, limited to shallow, turbid and highly productive habitat in the upper Gulf of California between Baja California and mainland Mexico (Rodriguez-Perez, Aurioles-Gamboa, Sanchez-Velasco, Lavin, & Newsome, 2018). The vaquita’s closest relatives are the congeneric Burmeister’s (*P. spinipinnis*) and spectacled (*P. dioptrica*) porpoises, which are found only in temperate and cold waters in the Southern Hemisphere, separated by at least 5000 km of ocean and two million years of divergence (Ben Chehida et al., in revision; McGowen, Spaulding, & Gatesy, 2009; Rosel, Haygood, & Perrin, 1995). Similar to other porpoises, vaquitas become entangled and die in gillnets set for finfish and shrimp (Rojas-Bracho & Reeves, 2013). The mortality rate was known to be unsustainable when studies on the bycatch rate (D’Agrosa, Lennert-Cody, & Vidal, 2000) and life history (Hohn, Read, Fernandez, Vidal, & Findley, 1996) were combined with the first abundance estimate of N=567 individuals (95% C.I. = 177-1073) in 1997 (Armando M. Jaramillo-Legorreta, Rojas-Bracho, & Gerrodette, 1999). The rate of decline has increased since approximately 2011 due to entanglement in illegal gillnets targeting totoaba (*Totoaba macdonaldi*), a large fish approximately the same size as the vaquita, captured for the black market trade of their swim bladders in China (Rojas-Bracho et al., 2019). The most recent estimates from 2018 indicate that fewer than 19 vaquita survive (A. M. Jaramillo-Legorreta et al., 2019). Initial genetics studies found no variation in mitochondrial DNA (mtDNA; Rosel & Rojas-Bracho, 1999) and low variation in the MHC DRB locus (Munguia-Vega et al., 2007). These authors have suggested that the low genetic diversity is due to long-term low effective population size (*Ne*) rather than to a recent bottleneck or the current rapid population decline (Munguia-Vega et al., 2007; Rojas-Bracho & Taylor, 1999; B. L. Taylor & Rojas-Bracho, 1999), but these data from few loci provide limited power to estimate timing or duration of demographic changes.

As part of the effort to prevent extinction of the vaquita and to further develop genomic resources to facilitate conservation and management planning for this and other endangered species, we used the Vertebrate Genomes Project (VGP) pipeline to generate a chromosomal-level, haplotype-phased reference vaquita genome assembly that exceeds the “platinum-quality” reference standards established by the VGP (Rhie et al., 2020a). The VGP standards are guidelines to ensure minimum error rates (QV40 or higher, or no more than 1 nucleotide error per 10,000 bp), highly contiguous and complete assemblies (contig N50 ≥ 1 Mb; chromosomal scaffold N50 ≥ 10 Mb), phasing of paternal and maternal haplotypes to reduce false gene duplication errors and manual curation to reduce errors and improve genome assembly quality. Based on the reference-quality assembly, we analyzed genomic diversity and historical demography to infer the cause of current low genomic diversity and whether genetic factors should be considered to be of concern for recovery if the immediate reason for decline, incidental bycatch in gillnets, can be halted in time to prevent extinction.

## Materials and Methods

### Genome data generation

Skin, mesovarium, kidney, trachea, and liver tissues were obtained during necropsy of the adult female vaquita that died during an attempt to begin ex-situ protection from illegal fishing operations (Rojas-Bracho et al., 2019). Cells were harvested and cultured at the Institute for Conservation Research, San Diego Zoo Global (Frozen Zoo®). From these cells, we generated a reference quality genome using the VGP pipeline 1.5 (Rhie et al., 2020a). In particular, we collected four genomic data types: Pacific Biosciences (Menlo Park, CA, USA) continuous long reads (CLR), 10X Genomics (Pleasanton, CA, USA) linked-reads, Bionano Genomics, Inc. (San Diego, CA, USA) DLS optical maps, and Arima Genomics, Inc. (San Diego, CA, USA) v1 Hi-C data. From one tube containing ~4 million cells in XPBS buffer with 10% DMSO and 10%

Glycerol, ultra-high molecular weight DNA (uHMW DNA) was extracted using the agarose plug Bionano Genomics protocol for Cell Culture DNA Isolation (Bionano Genomics, document No. 30026F). uHMW DNA quality was assessed by a Pulsed Field Gel assay and quantified with a Qubit 2 Fluorometer. From these extractions, 10 μg of uHMW DNA was sheared using a 26G blunt end needle (PacBio protocol PN 101-181-000 Version 05). A large-insert PacBio library was prepared using the Pacific Biosciences Express Template Prep Kit v1.0 (PN 101-357-000) following the manufacturer protocol. The library was then size selected (>20 kb) using the Sage Science BluePippin Size-Selection System and sequenced on 30 PacBio 1M v3 SMRT cells on the Sequel I instrument with the sequencing kit 3.0 (PN 101-597-800) and 10 hours movie. We used the same unfragmented DNA to generate a linked-reads library on the 10X Genomics Chromium linked-reads library (Genome Library Kit & Gel Bead Kit v2, PN 120258, Genome HT Library Kit & Gel Bead Kit v2, PN 120261, Genome Chip Kit v2, PN 120257, i7 Multiplex Kit, PN 120262). We sequenced this 10X Genomics library on an Illumina Novaseq S4 150 bp PE lane.

An aliquot of the same DNA was labeled for Bionano Genomics optical mapping using the Bionano Prep Direct Label and Stain (DLS) Protocol (document No. 30206E) and run on one Saphyr instrument chip flowcell. Hi-C reactions were performed by Arima Genomics according to the protocols described in the Arima-HiC kit (PN A510008). After the Arima-HiC protocol, Illumina-compatible sequencing libraries were prepared by first shearing purified Arima-HiC proximally-ligated DNA and then size-selecting DNA fragments from ~200-600 bp using SPRI beads. The size-selected fragments were then enriched for biotin and converted into Illumina-compatible sequencing libraries using the KAPA Hyper Prep kit (PN KK8504). After adapter ligation, DNA was PCR amplified and purified using SPRI beads. The purified DNA underwent standard QC (qPCR and Bioanalyzer (Agilent)) and was sequenced on the Illumina HiSeq X to ~60X coverage following the manufacturer’s protocols.

### Transcriptome data generation

Total RNA extraction and purification was conducted with QIAGEN RNAeasy kit (PN 74104). The quality and quantity of all RNAs were measured using a Fragment Analyzer (Aligent Technologies, Santa Clara, CA) and a Qubit 2.0 (Invitrogen). PacBio Iso-seq libraries were prepared according to the ‘Procedure & Checklist - Iso-Seq^™^ Template Preparation for Sequel^®^ Systems’ (PN 101-763-800 Version 01). Briefly, cDNA was reverse transcribed using the NEBNext^®^ Single Cell/Low Input cDNA Synthesis & Amplification Module (NEB E6421S) from 238 ng total RNA. Amplified cDNA was cleaned with 86 μl ProNex beads. The PacBio Iso-seq library was sequenced on one PacBio 8M (PN 101-389-001) SMRT Cell on the Sequel II instrument with sequencing kit 1.0 (PN 101-746-800) using the Sequel II Binding Kit 1.0 (PN 101-726-700) and 30 hours movie with two hours pre-extension.

The same RNA was used for mRNA-seq. The RNA-Seq library was prepared with 100 ng total RNA using the NEBNext Poly(A) mRNA Magnetic Isolation Module (NEB, PN E7490S) followed by NEBNext Ultra II Directional RNA Library Prep Kit for Illumina (PN E7760S). The library was then amplified over 14 cycles. Library quantification and qualification were performed with the Invitrogen Qubit dsDNA HS Assay Kit (PN Q32854). Libraries were sequenced on the Illumina NextSeq 500 in 150PE mid-output mode (Rockefeller Genomics Center). Data quality control was done using fastQC (v0.11.5; https://qubeshub.org/resources/fastqc).

### Genome assembly and annotation

We assembled the vaquita genome using the VGP 1.5 pipeline on the DNAnexus cloud computing system (https://platform.dnanexus.com/). Briefly, this pipeline is composed of an assembly step, scaffolding step and final polishing step. First, we assembled raw PacBio data with Falcon 2.0.0/Falcon-unzip 1.1.0 (Chin et al., 2016). Then, we polished the primary and alternate contigs using the same PacBio reads with arrow (PacBio smrtanalysis 6.0.0.47841). Prior to scaffolding, we detected and reassigned haplotype duplicated contigs in the primary contig set using purge_haplotig 1.0.4 (Roach, Schmidt, & Borneman, 2018) and we also extracted the mitochondrial reads to assemble the mitochondral sequence (Formenti et al., in prep). From this step, we only scaffolded the primary contigs using 10X Genomics data with scaff10x 4.1 (https://github.com/wtsi-hpag/Scaff10X), Bionano CMAP with Bionano Hybrid Solve 3.3_10252018 (Bionano Genomics) and Hi-C data with Salsa 2.2 (Ghurye, Pop, Koren, Bickhart, & Chin, 2017). Finally, the resulting primary scaffolds and alternate contigs were processed together through three polishing rounds: one additional round of arrow polishing and two rounds of polishing using 10X Illumina data mapped with Long Ranger 2.2.2 (https://github.com/10XGenomics/longranger) and base calling with FreeBayes 1.2.0 (Garrison & Marth, 2012). Primary scaffolds and alternate contigs were contamination checked and curated manually using gEVAL (Chow et al., 2016). For the primary assembly, this resulted in a further reduction of scaffold numbers by 11% and an increase of the scaffold N50 by 12% to 115 Mb. The primary and associated alternate assemblies were submitted to NCBI (accession GCA_008692025.1), and annotation was performed through their standard pipeline incorporating our RNA-seq and Iso-seq data (https://www.ncbi.nlm.nih.gov/genome/annotation_euk/process/). The primary assembly was screened for repetitive elements using RepeatMasker v4.0.5 (Smit, Hubley, & Green, 2013-2015) and the RepeatMasker combined database Dfam_Consensus-20181026. Base accuracy (QV) was measured using k=21 with Merqury (Rhie, Walenz, Koren, & Phillippy, 2020b). Gene content of the primary scaffolds was assessed using BUSCO v3.1.0 (Waterhouse et al., 2017) searches of the Laurasiatheria and mammalian gene set databases.

### Historical demography

To conduct analysis of historical demography using pairwise sequentially Markovian coalescent (PSMC; Li & Durbin, 2011), we first generated a diploid consensus genome from the 10X Genomics paired-end reads aligned to the primary haplotype assembly (Armstrong et al., 2019). The reads were trimmed with the BBduk function of BBTools (sourceforge.net/projects/bbmap/), removing the first 22 nucleotides of the R1 reads introduced during the Chromium library preparation (https://support.10xgenomics.com/genome-exome/library-prep/doc/technical-note-assay-scheme-and-configuration-of-chromium-genome-v2-libraries) and trimming all reads for average quality (q≥20), 3’ ends trimmed to q≥15 and minimum length (≥40 nucleotides). Unpaired reads were removed from the trimmed fastq files using the BBTools repair.sh function. Trimmed reads were aligned to the vaquita mitogenome (accession CM018178.1) using BWA mem (Li & Durbin, 2009), and the unmapped reads exported as reads representing only the nuclear genome. Nuclear reads were aligned to the primary haplotype assembly (accession GCA_008692025.1), and duplicate reads removed using Picard-Tools (http://broadinstitute.github.io/picard/). The resulting genome alignments from four 10X Genomics libraries were assessed for average depth of coverage using ANGSD (Korneliussen, Albrechtsen, & Nielsen, 2014), and combined for 47.8X average depth of coverage. From this coverage pile-up, the diploid consensus genome was extracted (Li & Durbin, 2011) and used as input for PSMC with generation time of 11.9 years based on the estimated generation time of harbor porpoise (Barbara L Taylor, Chivers, Larese, & Perrin, 2007), and an autosomal mutation rate (μA) of 1.08 x 10^-8^ substitutions per nucleotide per generation (Dornburg, Brandley, McGowen, & Near, 2012). PSMC atomic time intervals were combined as suggested by the authors (https://github.com/lh3/psmc) such that after 20 rounds of iterations, at least ~10 recombinations are inferred to occur in the intervals each parameter spans: p = (8+23*2+9+1). The remaining parameters were left as the default values used for humans (Li & Durbin, 2011), and we performed 100 bootstrap resamplings on all PSMC analyses to assess variance of the model.

### Genome-wide heterozygosity

The distribution of heterozygosity across the genome was determined using previously described analysis pipelines (Robinson et al., 2019). Briefly, we used HaplotypeCaller in the Genome Analysis Toolkit (GATK; McKenna et al., 2010) to call genotypes from the short-read pile-up (above), filtering out sites with <1/3X or >2X the average depth of coverage. Heterozygosity was calculated as the number of heterozygous sites divided by the total number of called genotypes in nonoverlapping 1Mb windows across each scaffold.

### Modeling demographic effects on heterozygosity

A coalescent simulation was constructed to estimate recent effective population size (*rN_e_*), historical effective population size (*hN_e_*) and time since a bottleneck (*b*) in which the population reduced in size from *hN_e_* to *rN_e_*. The analysis computed the likelihood of the empirical distribution of the number of heterozygous sites per kb (*H_kb_*) observed in 2244 1 Mb windows in the vaquita genome (from above) given similar distributions drawn from an equivalent genome arising from random draws of each of these parameters, which were sampled as:

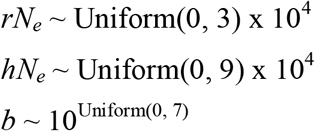

We initially drew 50,000 random values from these distributions. We then randomly selected 20,000 of these values where average growth rates (*(rN_e_* / *hN_e_*) / *b*) were less than 1.06, as values above this were considered to be biologically improbable (B. L. Taylor et al., 2019).

For each of the 20,000 scenarios, we generated one million independent SNPs for a single individual with a mutation rate of 1.08 x 10^-8^ substitutions/site/generation and a generation time of 11.9 years. To capture variability in the coalescent, we ran 4488 replicates of each scenario, which was twice the number of ~1 Mb windows in the empirical vaquita genome. This ensured that we could produce enough random sets of 2244 1 Mb windows from which to compute the scenario likelihoods as described below. The simulations were run with *fastsimcoal* v2.6.2 (Excoffier, Dupanloup, Huerta-Sanchez, Sousa, & Foll, 2013) through the R package *strataG* (v4.9.05).

For each of the 4488 replicates of one million SNPs in a scenario, we calculated the number of heterozygous SNPs per KB (*H’_kb_*). We then drew a random 2244 values of *H’_kb_* without replacement to represent one simulated genome for this scenario. We fit a gamma distribution to these values, which was used to compute the negative sum of log-likelihoods (−logL) of the empirical *Hkb* from the vaquita genome. For each scenario, we repeated this random draw of 2244 values of *H’_kb_* and computation of −logL 100 tiimes and recorded the mean and standard deviation of −logL. Likelihoods were plotted as heatmaps of the LOESS smoothed fit of −logL across pairs of simulation parameters. LOESS models were fit to each pair of parameters separately, and the surfaces represent the predicted −logL of 100,000 (10,000 x 10,000) evenly spaced points across each plot.

## Results

### A highly contiguous assembly of the vaquita genome

We assembled a 2.37 Gb genome (Table 1) in only 64 scaffolds, of which 21 represented arm-to-arm autosomes, named according to synteny with the blue whale (*Balaenoptera musculus*) and the X chromosome, in agreement with the 22-chromosome karyotype. The remaining 42 unplaced scaffolds consisted of only 0.198 Gb combined (0.08% of the total length), meaning that 99.92% of the assembled sequence has been assigned to chromosomes. Consistent with this mostly complete assembly, the N50 contig value was 20.22 Mb (273 contigs), N50 scaffold was 115.47 Mb, and base call accuracy was QV40.88 (0.82 errors per 10,000 bp). There were only 208 gaps, of which the annotated chromosomes had 3-17 gaps each. The Hi-C heat-map showing genomic interactions (Figure 1) indicates strong agreement between the close interactions and chromosome-length scaffolds. The alternate haplotype contigs are made up of 1 Gb of the genome, indicating low heterozygosity. Depth of coverage for each data type are presented in Table 2. Assemblies of both primary and alternate haplotypes have been deposited at DDBJ/ENA/GenBank under the accessions VOSU00000000 (principle haplotype) and VOSV00000000 (alternate haplotype) in BioProjects PRJNA557831 and PRJNA557832, respectively.

**Figure 1.**
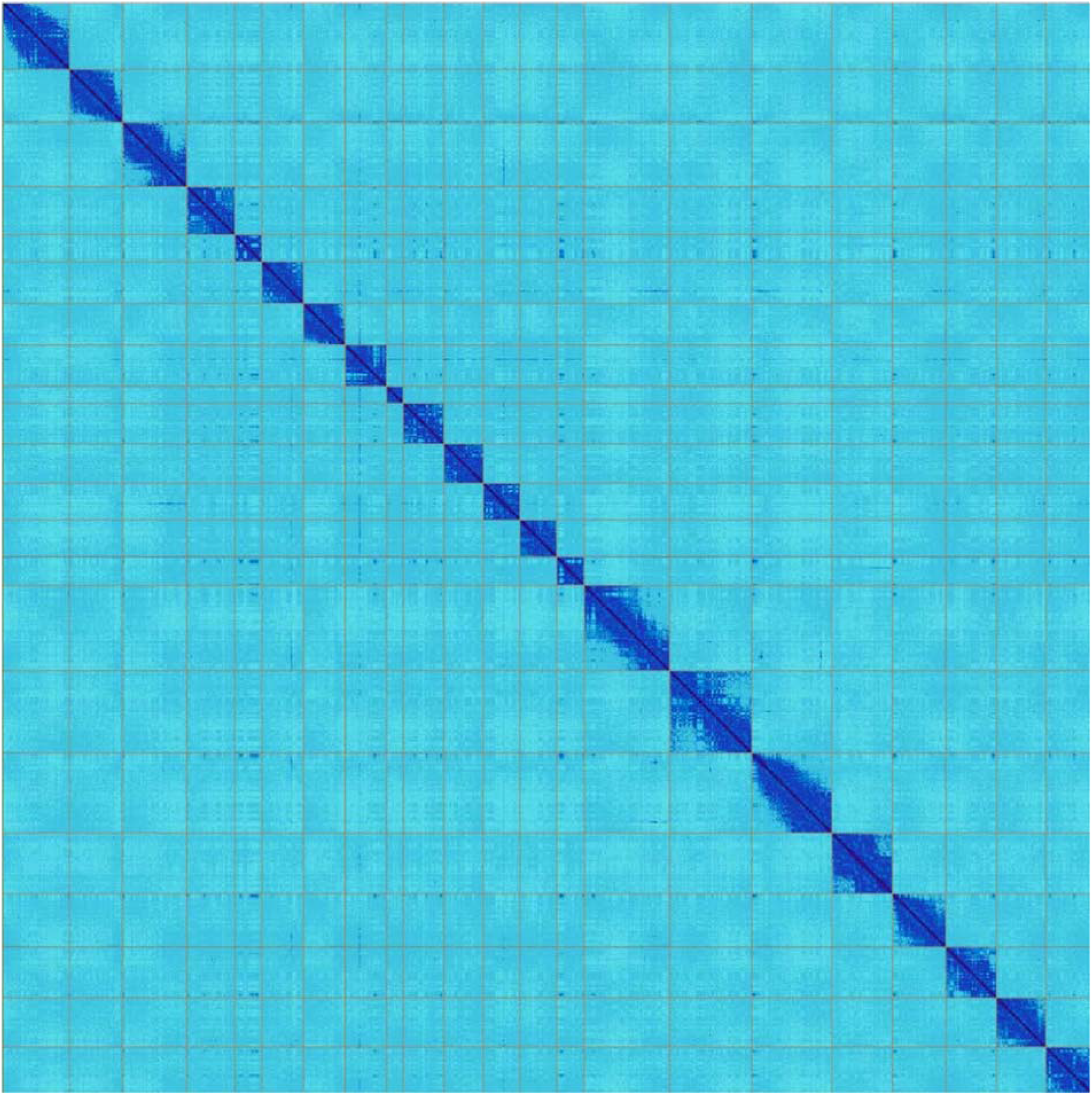
HiC heat-map of genomic interactions. Interactions between two locations are depicted by a dark blue pixel. Gray lines depict scaffold boundaries for the 22 chromosome-length scaffolds. Different scaffolds should not share any interactions (pixels off diagonal outside the scaffold boundaries), while patterns within a scaffold show chromosome-substructure.

**Table 1.**
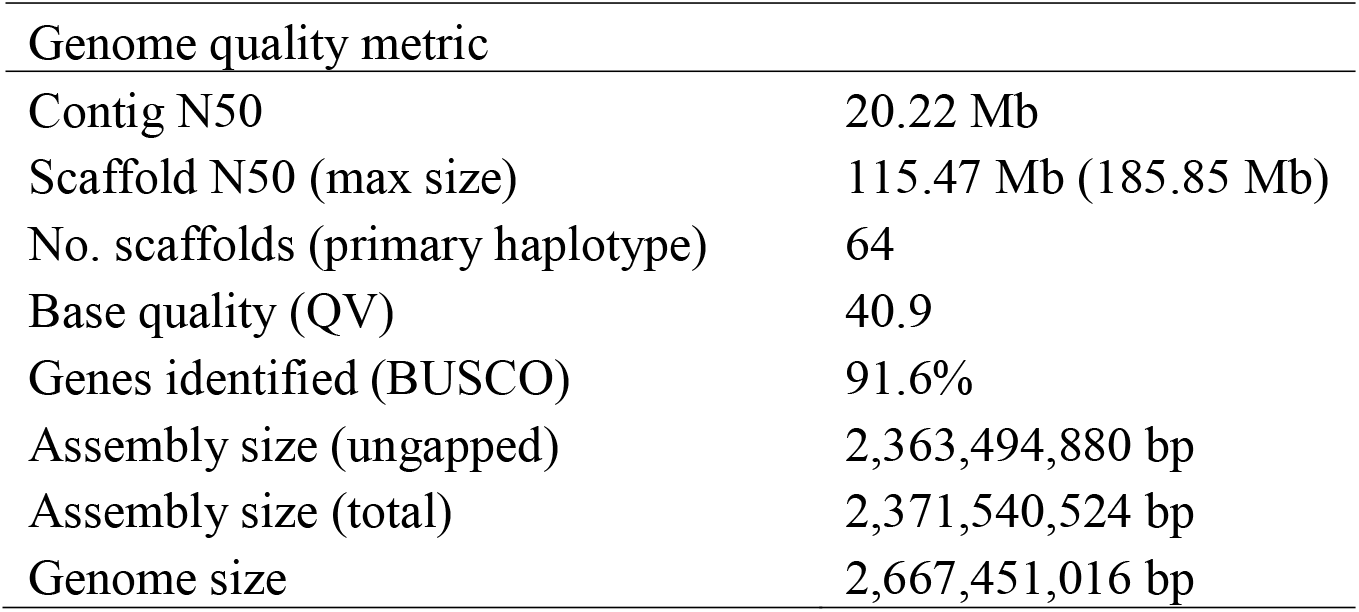
Vaquita genome assembly metrics. Genome size is the kmer estimate based on GenomeScope (v1.0) analysis of the 10X Genomics data with k = 31. The BUSCO score is for complete genes identified from the mammalian single-copy conserved gene data set.

| Genome quality metric |  |
| --- | --- |
| Contig N50 | 20.22 Mb |
| Scaffold N50 (max size) | 115.47 Mb (185.85 Mb) |
| No. scaffolds (primary haplotype) | 64 |
| Base quality (QV) | 40.9 |
| Genes identified (BUSCO) | 91.6% |
| Assembly size (ungapped) | 2,363,494,880 bp |
| Assembly size (total) | 2,371,540,524 bp |
| Genome size | 2,667,451,016 bp |

**Table 2.** Estimated genome sequence average depth of raw data coverage (before adapter and quality trimming) for sequencing and mapping technologies based on an estimated genome size of 2.7 Gb.

| Data type | Raw data (bp) | Coverage |
| --- | --- | --- |
| 10x Genomics | 200,218,960,380 | 74X |
| Arima Genomics HiC | 255,724,383,000 | 94X |
| Bionano Genomics | 480,155,600,000 | 178X |
| PacBio SubReads | 325,960,000,000 | 121X |

BUSCO analysis showed 89.9% and 91.6% gene content identification from the primary haplotype when compared to the Laurasiatheria and mammalian data sets, respectively, with only 1.0 and 1.1% of the complete genes duplicated, respectively, and 4.3 and 4.6% fragmented (Supplemental Table S1). Genome annotation identified 26,497 genes and pseudogenes, 19,069 of which are protein coding (Table S2). The cumulative number of genes with alignment to the UniProtKB/Swiss-Prot curated proteins was 18,748 (89%) at ≥90% coverage of the target protein. This coverage was 5-48% higher than the number of genes aligned from other annotated cetacean genomes (Table S2). Similar to other cetacean genomes (e.g., Fan et al., 2019; Keane et al., 2015; Tollis et al., 2019), the vaquita genome consisted of about 46% repeats (Table 3) based on RepeatMasker.

**Table 3.** Repetitive content of the vaquita genome (total assembly length 23.72 Gb) as determined by RepeatMasker.

| Repeat type | Length (bp) | % of Genome |
| --- | --- | --- |
| SINEs | 189,109,608 | 7.97% |
| LINEs | 653,546,597 | 27.56% |
| LTR | 134,757,334 | 5.68% |
| DNA transposons | 76,591,695 | 3.23% |
| Unclassified | 1,047,864 | 0.04% |
| Satellites | 1,588,863 | 0.07% |
| Simple repeats | 23,753,228 | 1% |
| Low complexity | 4,527,734 | 0.19% |
| Total repeats: | 1,085,270,145 | 45.76% |

### Low heterozygosity of the vaquita genome

Genome-wide heterozygosity was 0.0105% overall, with even distribution of heterozygosity across the genome (Figure 2A). Heterozygosity per 1 Mb window ranged from 0 to 1.2/kb, but only two (noncontiguous) windows out of 2247 had no heterozygotes, and the standard deviation of heterozygosity across the windows was very low (SD = 0.0000767). None of the 1 Mb windows had heterozygosity of >1.3/kb, and 94% of the windows had heterozygosity of <0.2/kb (Figure 2B). In comparison to other mammals, the vaquita genome exhibits the lowest heterozygosity yet detected in an outbreeding mammalian species (Figure 3), with the exception of the San Nicolas Island fox (*Urocyon littoralis*), an endemic subspecies found only on a 58 km^2^ island approximately 100 km off the coast of California, with an estimated population size of about 500 individuals (Robinson et al., 2016). However, unlike the vaquita, heterozygosity is not evenly distributed across the genome in the San Nicolas Island fox and other small inbred populations of canids, due to the effects of recent inbreeding in addition to long-term small population sizes (Robinson et al., 2019).

**Figure 2.**
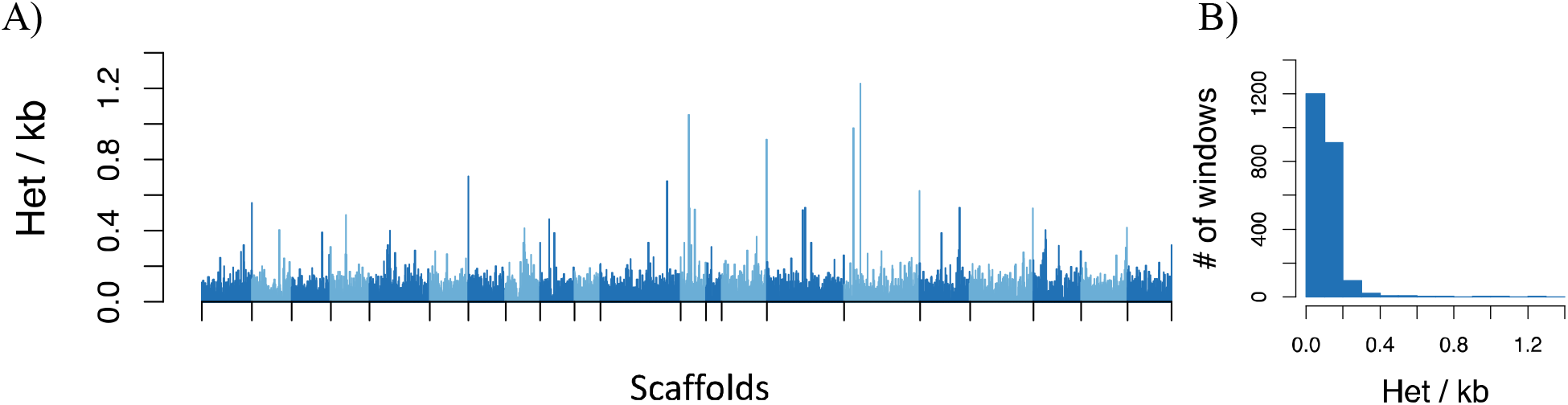
Distributions of heterozygosity across the vaquita genome. A) Bar plot shows persite heterozygosity in nonoverlapping 1-Mb windows across 22 scaffolds >10 Mb in length. Scaffolds are shown in alternating shades. B) Histogram of the count of per-window heterozygosity levels.

**Figure 3.**
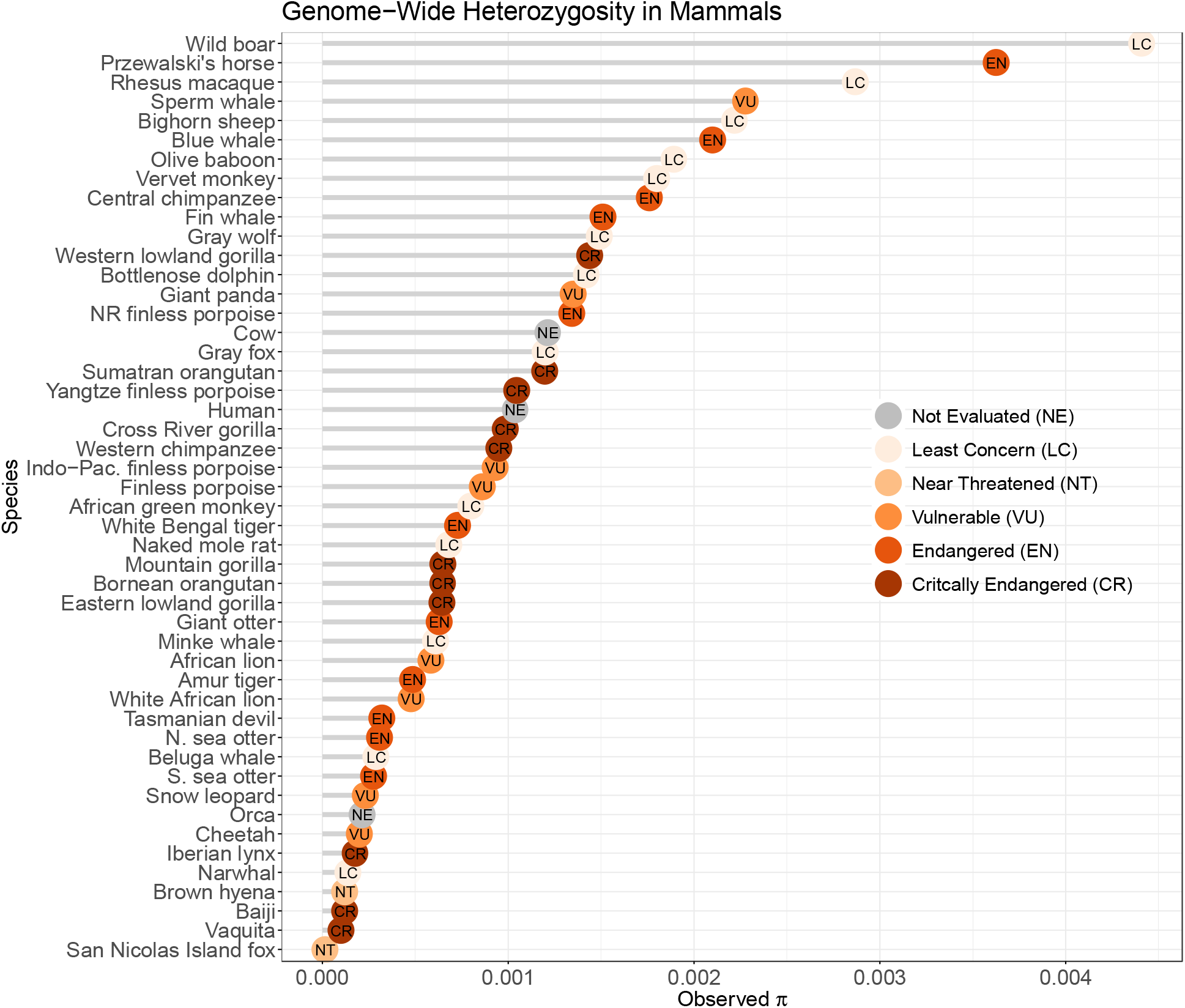
Comparison of genome-wide heterozygosity (π) among mammals. Values are drawn from the literature, based on Robinson et al. (2016), plus the vaquita and blue whale. Dots are colored by the endangered status according to the Red List for Threatened Species, International Union for Conservation of Nature (IUCN). Although the Baiji, or Yangtze River dolphin, is listed as critically endangered, it is believed to have been extinct since at least 2006 (Turvey et al., 2007). See supplemental Table S4 for heterozygosity information.

### Vaquita population size over time

This low, relatively even heterozygosity across the vaquita genome could be indicative of a long-term small, outbred population (Robinson et al., 2019; Westbury et al., 2019) To test this hypotheses, we performed PSMC analysis. The results indicates that the vaquita effective population size has been small, ranging from about 1,400 to 3,200 for most of the last ~300,000 years (Figure 4A). This finding corroborates previous conclusions based on single-locus analyses (Munguia-Vega et al., 2007; B. L. Taylor & Rojas-Bracho, 1999) but extends the duration of persistence of the species at low *Ne* to the mid Pleistocene, prior to the penultimate glacial period, the Saalian, which lasted from approximately 300,000 to 130,000 years ago.

**Figure 4.**
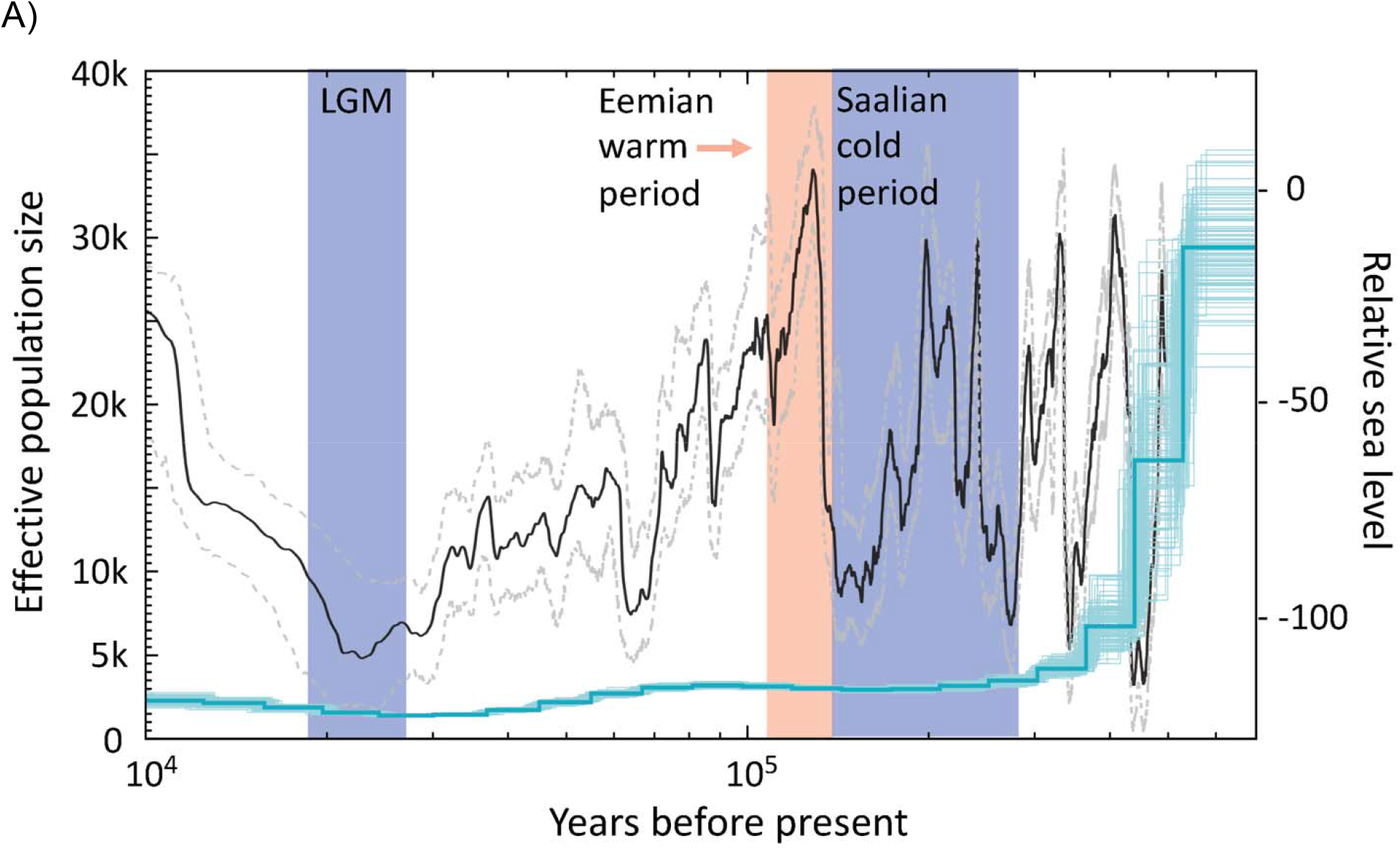
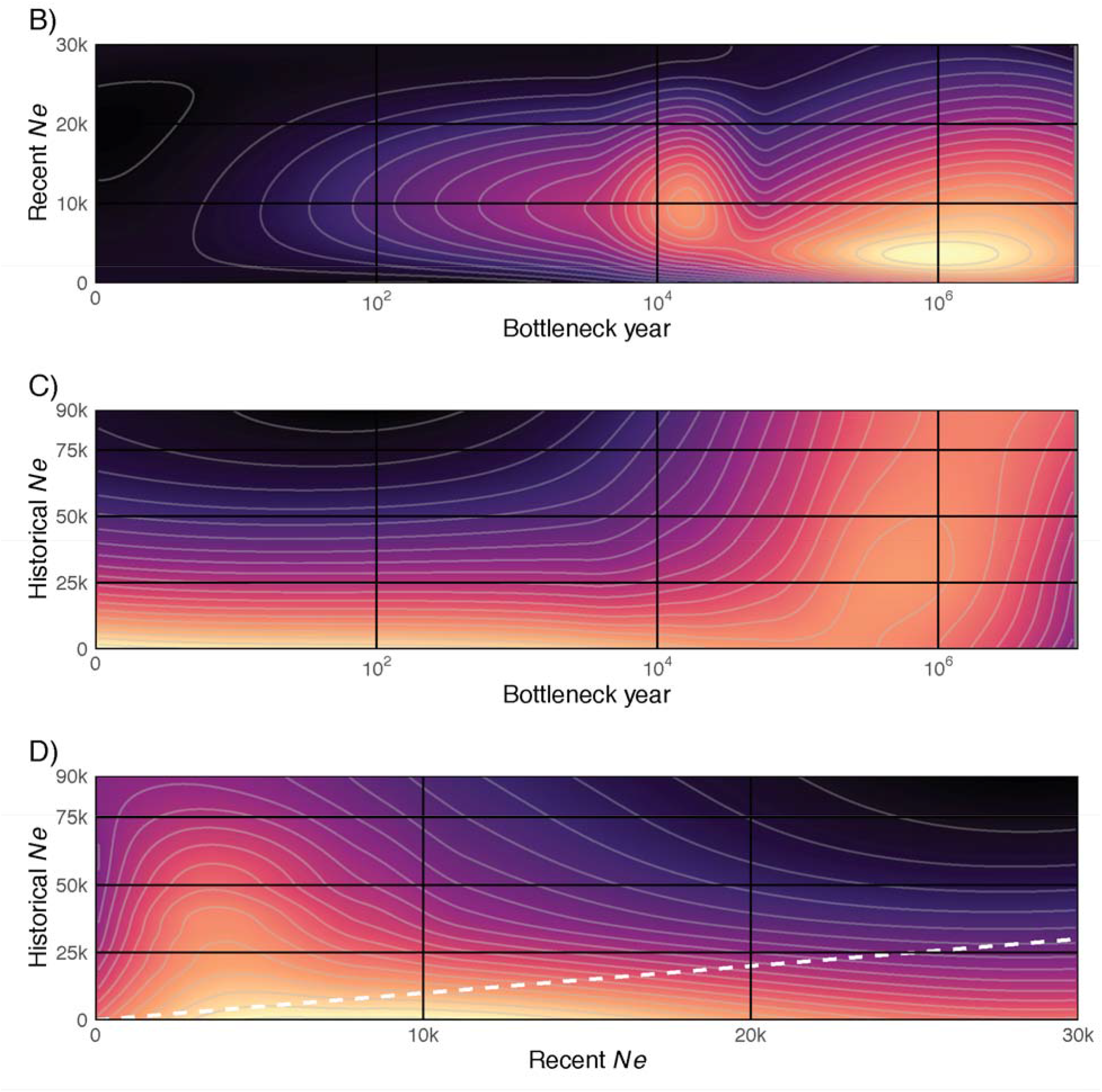
Changes in vaquita population size over time. A) Changes in effective population size (*Ne*) of the vaquita over time inferred from PSMC analysis of the nuclear genome. The darker blue line represents the median and lighter lines represent the 100 bootstrap replicates. The black line shows relative sea level (right axis, compared to present) with 95% confidence intervals (gray dashed lines) from Grant et al. (2014), and shading corresponds to cold and warm periods. B – D) Heatmap of the distribution of the negative log-likelihood (−logL) of the empirical heterozygosity distribution across pairs of demographic parameters from the coalescent model, with higher likelihood combinations shown by lighter color. The dashed white line in (D) represents a 1:1 slope, where current and historical population sizes would have been equal before and after the modeled change in population size.

## Discussion

We have assembled the most complete cetacean genome to date, as measured by the low number of scaffolds, small number of gaps per chromosome scaffold, high percentage of scaffolds assigned to 22 chromosomes, cumulative number of genes with an alignment to the UniProtKB/Swiss-Prot curated proteins and small amount of missing data. Identification of gene content was also in the expected range for a high-quality mammalian genome at 90.5% of complete single-copy genes from the BUSCO mammalian gene set, with a low level of false duplicates and low levels of fragmented genes.

The PSMC analysis indicates that the vaquita population declined during the late Pleistocene, most likely due to climate change and the associated habitat changes in the eastern North Pacific coastal regions of North and Central America, and that it remained small over the last approximately 300,000 years. PSMC results can be affected by population structure, inbreeding, changes in connectivity among populations and stochastic variation in coalescent events when diversity is low (Beichman, Phung, & Lohmueller, 2017; Li & Durbin, 2011; Mazet, Rodriguez, Grusea, Boitard, & Chikhi, 2016; Orozco-terWengel, 2016). The coalescent results are consistent with the PSMC-inferred historical demography being the most likely cause of current heterozygosity levels rather than a recent severe bottleneck or inbreeding. Importantly, the duration of the small population size indicates that the observed level of heterozygosity is the result of a population at genetic equilibrium, where mutations are balanced by drift and selection, and that highly deleterious mutations are likely to have been purged from the population (Day, Bryant, & Meffert, 2003; Dussex et al., in revision; Robinson et al., 2018; Westbury et al., 2018; Westbury et al., 2019).

Examples of species with low diversity but long-term viability and potential for adaptability are becoming more common (Dussex et al., in revision; Andrew D Foote et al., 2019; Robinson et al., 2018; Westbury et al., 2018; Westbury et al., 2019; Xue et al., 2015). Among odontocetes (toothed whales, dolphins and porpoises), in particular, there are examples of species with nearly as low diversity as the vaquita that exhibit strong evidence of the influence of demographic factors influencing genome-wide diversity over tens to hundreds of thousands of years of diversification and adaptation (Andrew D Foote et al., 2019; Andrew D Foote et al., 2016; Van Cise et al., 2019; Westbury et al., 2019). In several of these cases where it has been examined, genome-wide heterozygosity patterns do not indicate that low diversity was caused by rapid bottlenecks or inbreeding; instead, these patterns indicate that low diversity has been present for extended periods while species persist and diversify (e.g., narwhal (Westbury et al., 2019), orca (Andrew D Foote et al., 2019)). These examples and others (Robinson et al., 2018; Robinson et al., 2016; Westbury et al., 2018) indicate that, contrary to the paradigm of an “extinction vortex” (Gilpin & Soulé, 1986) that may doom species with low diversity, some species have persisted with low genomic diversity and small population size. Long-term small population size enables the purging of recessive deleterious alleles, thereby reducing the risk of inbreeding depression, perhaps allowing for continued future persistence with relatively small population sizes and an increased tolerance to the genetic consequences of bottlenecks.

The vaquita’s current habitat in the upper Gulf of California was likely diminished or absent due to low sea levels several times through the last 350,000 years (Siddall et al., 2003), with the lowest sea level occurring at the end of the Saalian complex and the LGM (Figure 2) followed by a rapid rise of 120-140 m (similar to the present level) during the Eemian warm period between 115,000 and 130,000 years ago and after the LGM (Figure 5). Over much of the last 100,000 years, sea level has been intermediate between the high points (present and Eemian warm period) and lows (end of Saalian and the LGM) (Rohling et al., 2017). There is no fossil record or other indication that vaquita have ever inhabited colder parts of the eastern North Pacific along the west coast of Baja California, Mexico, or further north off of California at the southern end of the current range of the congeneric harbor porpoise (*Phocoena phocoena*) (Brownell Jr., 1983). The closest relative of the vaquita, the Burmeister’s porpoise or the ancestor of two sister species, Burmeister’s and spectacled porpoise (Ben Chehida et al., in revision), are both found only in temperate and cold waters of the southern hemisphere. Based on the closer relationship to southern hemisphere species and on the similar timing of rapid climate warming and vaquita population decline, it appears that climate change at the end of the Saalian ice age caused a northward shift of the species range, resulting in a remnant population being isolated in the Gulf of California, where it has persisted in the newly expanded and shallow, highly productive upper Gulf region.

**Figure 5.**
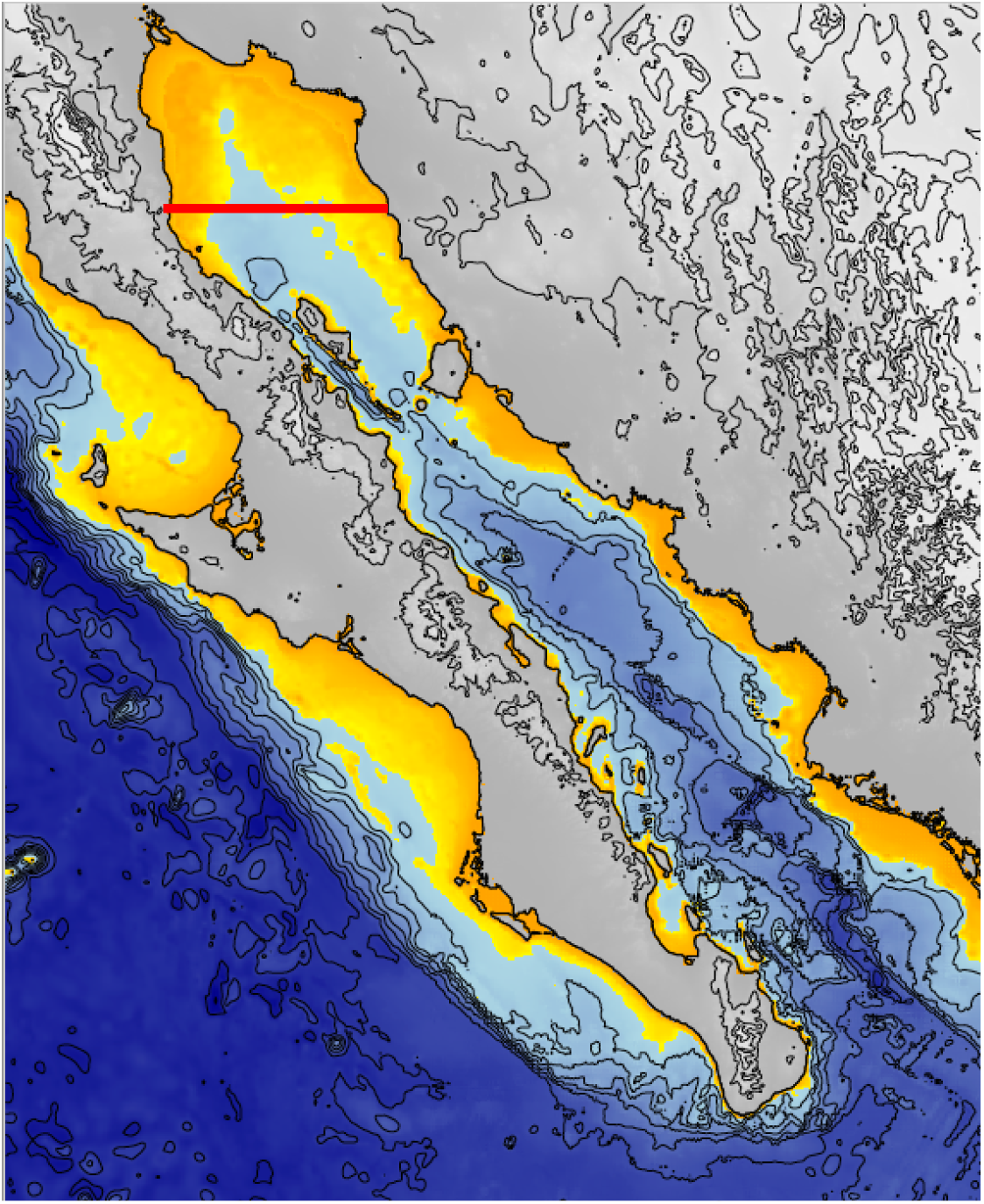
Bathymetric map of the Gulf of California showing 500m isobath lines. Transition to yellow is at −140m, indicating portions of the Gulf that were likely above sea level during the last two glacial maxima, ~22,000 and 140,000 years ago. The area north of the red line is the approximate historical range of the vaquita (Brownell, 1986).

The reference genome presented here has provided important insight into the demographic history of the critically endangered vaquita, reinforcing a previous hypothesis (B. L. Taylor & Rojas-Bracho, 1999) that the low genetic diversity of the vaquita is not due to a recent extreme bottleneck or current inbreeding. These results taken together with recent evidence of healthy looking vaquitas, often with robust calves (B. L. Taylor et al., 2019), suggest that population recovery may not be hindered because of genetic issues. Analysis of re-sequenced genomes from multiple individuals sampled over the previous few decades will shed light on changes in inbreeding as the population has declined due to bycatch in gillnets, and whether deleterious mutations are likely to have been purged from the genome as a result of the long-term persistence at a small population size, as has been suggested for some other species and populations (e.g., Dussex et al., in revision; Robinson et al., 2018; Westbury et al., 2018; Westbury et al., 2019).

Finally, this genome assembly is the highest quality, most complete genome in the odontocete lineage that consists of all dolphins, porpoises and toothed whales. As such, it provides a genomic resource for better reference-guided assemblies and scaffolding of other cetacean genomes (Alonge et al., 2019; Lischer & Shimizu, 2017; Morin et al., in revision) and for comparative genomics, especially for variation in genome structure. We expect that the vaquita genome, along with expected assembly of reference genomes for other endangered species, will continue to contribute to both understanding and conservation of global biodiversity (Kraus et al., submitted).

## Acknowledgements

The planning and diligence of the VaquitaCPR team were critical in collection, preservation and rapid delivery of tissue samples for tissue culture that made this project possible. We are grateful to all people involved in obtaining and culturing the tissue samples. Kelly Robertson was instrumental in ensuring rapid import of the samples under CITES permit (permit No. 17US774233/9; Mexican export permit MX89760). Tissue samples are stored in the SWFSC Marine Mammal and Sea Turtle Research (MMASTR) collection under MMPA permit 19091. We thank Annabel Beichman for help with PSMC analysis and Prof. Eelco Rohling for assistance with interpretation of sea level data. We are grateful to Cisco Werner, Director of Scientific Programs and Chief Science Advisor for NOAA Fisheries, for funding sequencing of the vaquita genome. Earlier versions of the manuscript have been improved thanks to careful review by Love Dalén and Mark Chaisson.

## Data Availability

The vaquita reference genome and all sequence data are available via the Vertebrate Genome Project GenomeArk website (https://vgp.github.io/genomeark/Phocoena_sinus/) and NCBI Genome database (Bioprojects PRJNA557831 and PRJNA557832). Annotation is available at NCBI (http://www.ncbi.nlm.nih.gov/genome/annotation_euk/Phocoena_sinus/100/). Ensembl annotation for the vaquita is available via the VGP pre-release data portal (projects.ensembl.org/vgp) and will be fully integrated into the Ensembl genome browser (ensembl.org), including comparative data, in release 101, due to go live by August 2020.

## Author Contributions

PAM, EDJ and OAR initiated the project, and PAM, EDJ and OF designed and led research and analyses and co-wrote the manuscript. FIA, BH, JRB, JM, and OF generated data. JM and SPaez initiated the project for the VGP. AP, AR, BH, AF, GF, KH, JR, JTorrence, MJPC, WC, SPalen and YVB contributed to data processing and genome assembly. MLH, ACM, JAF and CDA cultured cell lines, and AW, BLT, CRS, FMDG, JTeilmann, LR-B, MPH-J, RSW, SS, TR and WM conducted the field work to obtain and process the tissue samples. All authors contributed to interpretation of results and preparation of the manuscript.

## References

Alonge, M., Soyk, S., Ramakrishnan, S., Wang, X., Goodwin, S., Sedlazeck, F. J., … Schatz, M. C. (2019). RaGOO: fast and accurate reference-guided scaffolding of draft genomes. Genome Biology, 20(1), 224. doi:10.1186/s13059-019-1829-6

Armstrong, E. E., Taylor, R. W., Prost, S., Blinston, P., van der Meer, E., Madzikanda, H., … Petrov, D. (2019). Cost-effective assembly of the African wild dog (*Lycaon pictus*) genome using linked reads. Gigascience, 8(2). doi:10.1093/gigascience/giy124

Arnason, U., Lammers, F., Kumar, V., Nilsson, M. A., & Janke, A. (2018). Whole-genome sequencing of the blue whale and other rorquals finds signatures for introgressive gene flow. Science Advances, 4(4), eaap9873. doi:10.1126/sciadv.aap9873

Autenrieth, M., Hartmann, S., Lah, L., Roos, A., Dennis, A. B., & Tiedemann, R. (2018). High-quality whole-genome sequence of an abundant Holarctic odontocete, the harbour porpoise (*Phocoena phocoena*). Molecular Ecology Resources, 18(6), 1469–1481. doi:10.1111/1755-0998.12932

Beichman, A. C., Phung, T. N., & Lohmueller, K. E. (2017). Comparison of single genome and allele frequency data reveals discordant demographic histories. G3, 7(11), 3605–3620. doi:10.1534/g3.117.300259

Ben Chehida, Y., Aguilar, A., Borrel, A., Ferreira, M., Taylor, B. L., Rojas-Bracho, L., … Fontaine, M. C. (in revision). Evolutionary history of the porpoise family (Phocoenidae) across the speciation continuum: a phylogeographic perspective from the complete mitochondrial genome. BioRxiv doi 10.1101/851469. doi:10.1101/851469

Brownell Jr., R. L. (1983). Phocoena sinus. Mammalian Species, 198, 1–3.

Chattopadhyay, B., Garg, K. M., Yun Jing, S., Low, G. W., Frechette, J., & Rheindt, F. E. (2019). Conservation genomics in the fight to help the recovery of the critically endangered Siamese crocodile *Crocodylus siamensis*. Molecular Ecology, 28(5), 936–950. doi:10.1111/mec.15023

Chin, C. S., Peluso, P., Sedlazeck, F. J., Nattestad, M., Concepcion, G. T., Clum, A., … Schatz, M. C. (2016). Phased diploid genome assembly with single-molecule real-time sequencing. Nature Methods, 13(12), 1050–1054. doi:10.1038/nmeth.4035

Chow, W., Brugger, K., Caccamo, M., Sealy, I., Torrance, J., & Howe, K. (2016). gEVAL - α web-based browser for evaluating genome assemblies. Bioinformatics, 32(16), 2508–2510. doi:10.1093/bioinformatics/btw159

D’Agrosa, C., Lennert-Cody, C. E., & Vidal, O. (2000). Vaquita bycatch in Mexico’s artisanal gillnet fisheries: Driving a small population to extinction. Conservation Biology, 14(4), 1110–1119. doi:10.1046/j.1523-1739.2000.98191.x

Day, S. B., Bryant, E. H., & Meffert, L. M. (2003). The influence of variable rates of inbreeding on fitness, environmental responsiveness, and evolutionary potential. Evolution, 57(6), 1314–1324. doi:10.1111/j.0014-3820.2003.tb00339.x

Dornburg, A., Brandley, M. C., McGowen, M. R., & Near, T. J. (2012). Relaxed clocks and inferences of heterogeneous patterns of nucleotide substitution and divergence time estimates across whales and dolphins (Mammalia: Cetacea). Molecular Biology and Evolution, 29(2), 721–736. doi:10.1093/molbev/msr228

Dussex, N., van der Valk, T., Wheat, C. W., Diez-del-Molino, D., von Seth, J., Foster, Y., … Dalén, L. (in revision). Population genomics reveals the impact of small population size in the critically endangered kākāpō Nature Ecology and Evolution.

Excoffier, L., Dupanloup, I., Huerta-Sanchez, E., Sousa, V. C., & Foll, M. (2013). Robust demographic inference from genomic and SNP data. PLoS Genetics, 9(10), e1003905. doi:10.1371/journal.pgen.1003905

Fan, G., Zhang, Y., Liu, X., Wang, J., Sun, Z., Sun, S., … Liu, X. (2019). The first chromosome-level genome for a marine mammal as a resource to study ecology and evolution. Molecular Ecology Resources, 19(4), 944–956. doi:10.1111/1755-0998.13003

Foote, A. D., Liu, Y., Thomas, G. W., Vinar, T., Alfoldi, J., Deng, J., … Gibbs, R. A. (2015). Convergent evolution of the genomes of marine mammals. Nature Genetics, 47(3), 272–275. doi:10.1038/ng.3198

Foote, A. D., Martin, M. D., Louis, M., Pacheco, G., Robertson, K. M., Sinding, M.-H. S., … Morin, P. A. (2019). Killer whale genomes reveal a complex history of recurrent admixture and vicariance. Molecular Ecology, 28, 3427–3444. doi:10.1111/mec.15099

Foote, A. D., Vijay, N., Ávila-Arcos, M. C., Baird, R. W., Durban, J. W., Fumagalli, M., … Wolf, J. B. W. (2016). Genome-culture coevolution promotes rapid divergence in the killer whale. Nature Communications, 7, Article No. 11693. doi:10.1038/ncomms11693

Formenti, G., Fedrigo, O., Howe, K., Balacco, J., Haase, B., Mountcastle, J., … Jarvis, E. D. (in prep). Complete and near error-free mitochondrial genome assemblies with long reads reveal unreported gene duplications and repeats. Nature.

Garner, B. A., Hand, B. K., Amish, S. J., Bernatchez, L., Foster, J. T., Miller, K. M., … Luikart, G. (2016). Genomics in conservation: case studies and bridging the gap between data and application. Trends in Ecology and Evolution, 31(2), 81–83. doi:10.1016/j.tree.2015.10.009

Garrison, E., & Marth, G. (2012). Haplotype-based variant detection from short-read sequencing. arXiv:1207.3907[q-bio.GN].

Ghurye, J., Pop, M., Koren, S., Bickhart, D., & Chin, C. S. (2017). Scaffolding of long read assemblies using long range contact information. BMC Genomics, 18(1), 527. doi:10.1186/s12864-017-3879-z

Gilpin, M. E., & Soulé, M. E. (1986). Minimum viable populations: processes of species extinction. In M. E. Soulé (Ed.), Conservation Biology: The science of scarcity and diversity (pp. 19–34): Sinauer.

Harrisson, K. A., Pavlova, A., Telonis-Scott, M., & Sunnucks, P. (2014). Using genomics to characterize evolutionary potential for conservation of wild populations. Evolutionary Applications, 7(9), 1008–1025. doi:10.1111/eva.12149

He, X., Johansson, M. L., & Heath, D. D. (2016). Role of genomics and transcriptomics in selection of reintroduction source populations. Conservation Biology, 30(5), 1010–1018. doi:10.1111/cobi.12674

Hedrick, P. W., Robinson, J. A., Peterson, R. O., & Vucetich, J. A. (2019). Genetics and extinction and the example of Isle Royale wolves. Animal Conservation, 22(3), 302–309. doi:10.1111/acv.12479

Hohn, A. A., Read, A. J., Fernandez, S., Vidal, O., & Findley, L. T. (1996). Life history of the vaquita, *Phocoena sinus* (Phocoenidae, Cetacea). Journal of Zoology, 239, 235–251. doi:DOI 10.1111/j.1469-7998.1996.tb05450.x

Jaramillo-Legorreta, A. M., Cardenas-Hinojosa, G., Nieto-Garcia, E., Rojas-Bracho, L., Thomas, L., Ver Hoef, J. M., … Tregenza, N. (2019). Decline towards extinction of Mexico’s vaquita porpoise (*Phocoena sinus*). Royal Society Open Science, 6(7). doi:10.1098/rsos.190598

Jaramillo-Legorreta, A. M., Rojas-Bracho, L., & Gerrodette, T. (1999). A new abundance estimate for vaquitas: First step for recovery. Marine Mammal Science, 15(4), 957–973. doi:10.1111/j.1748-7692.1999.tb00872.x

Keane, M., Semeiks, J., Webb, A. E., Li, Y. I., Quesada, V., Craig, T., … de Magalhaes, J. P. (2015). Insights into the evolution of longevity from the bowhead whale genome. Cell Reports, 10(1), 112–122. doi:10.1016/j.celrep.2014.12.008

Korneliussen, T. S., Albrechtsen, A., & Nielsen, R. (2014). ANGSD: Analysis of next generation sequencing data. BMC Bioinformatics, 15, 356. doi:10.1186/s12859-014-0356-4

Kraus, R. H. S., Paez, S., Ceballos, G., Crawford, A., Fedrigo, O., Finnegan, S., … Jarvis, E. D. (submitted). The role of genomics in conserving biodiversity during the sixth mass extinction. Nature Reviews Genetics.

Li, H., & Durbin, R. (2009). Fast and accurate short read alignment with Burrows-Wheeler transform. Bioinformatics, 25(14), 1754–1760. doi:10.1093/bioinformatics/btp324

Li, H., & Durbin, R. (2011). Inference of human population history from individual whole-genome sequences. Nature, 475(7357), 493–496. doi:10.1038/nature10231

Lischer, H. E. L., & Shimizu, K. K. (2017). Reference-guided de novo assembly approach improves genome reconstruction for related species. BMC Bioinformatics, 18(1), 474. doi:10.1186/s12859-017-1911-6

Mazet, O., Rodriguez, W., Grusea, S., Boitard, S., & Chikhi, L. (2016). On the importance of being structured: instantaneous coalescence rates and human evolution--lessons for ancestral population size inference? Heredity (Edinb), 116(4), 362–371. doi:10.1038/hdy.2015.104

McGowen, M. R., Spaulding, M., & Gatesy, J. (2009). Divergence date estimation and a comprehensive molecular tree of extant cetaceans. Molecular Phylogenetics and Evolution, 53(3), 891–906. doi:10.1016/j.ympev.2009.08.018

McKenna, A., Hanna, M., Banks, E., Sivachenko, A., Cibulskis, K., Kernytsky, A., … DePristo, M. A. (2010). The Genome Analysis Toolkit: a MapReduce framework for analyzing next-generation DNA sequencing data. Genome Research, 20(9), 1297–1303. doi:10.1101/gr.107524.110

Morin, P. A., Alexander, A., Blaxter, M., Caballero, S., Fedrigo, O., Fontaine, M. C., … Jarvis, E. D. (in revision). Building genomic infrastructure: Sequencing platinum-standard reference-quality genomes of all cetacean species. Marine Mammal Science.

Morin, P. A., Foote, A. D., Baker, C. S., Hancock-Hanser, B. L., Kaschner, K., Mate, B. R., … Alexander, A. (2018a). Demography or selection on linked cultural traits or genes? Investigating the driver of low mtDNA diversity in the sperm whale using complementary mitochondrial and nuclear genome analyses. Molecular Ecology, 27(11), 2604–2619. doi:10.1111/mec.14698

Morin, P. A., Foote, A. D., Hill, C. M., Simon-Bouhet, B., Lang, A. R., & Louise, M. (2018b). SNP discovery from single and multiplex genome assemblies of non-model organisms. In S. R. Head, P. Ordoukhanian, & D. Salomon (Eds.), Next-Generation Sequencing. Methods in Molecular Biology (Vol. 1712, pp. 113–144): Humana Press.

Munguia-Vega, A., Esquer-Garrigos, Y., Rojas-Bracho, L., Vazquez-Juarez, R., Castro-Prieto, A., & Flores-Ramirez, S. (2007). Genetic drift vs. natural selection in a long-term small isolated population: major histocompatibility complex class II variation in the Gulf of California endemic porpoise (*Phocoena sinus*). Molecular Ecology, 16(19), 4051–4065. doi:10.1111/j.1365-294X.2007.03319.x

Norris, K. S., & McFarland, W. N. (1958). A new harbor porpoise of the genus *Phocoena* from the Gulf of California. Journal of Mammalogy, 39, 22–39.

Orozco-terWengel, P. (2016). The devil is in the details: the effect of population structure on demographic inference. Heredity, 116(4), 349–350. doi:10.1038/hdy.2016.9

Rhie, A., McCarthy, S. A., Fedrigo, O., Damas, J., Formenti, G., Koren, S., … Jarvis, E. D. (2020a). Towards complete and error-free genome assemblies of all vertebrate species. bioRxiv. doi:10.1101/2020.05.22.110833

Rhie, A., Walenz, B. P., Koren, S., & Phillippy, A. M. (2020b). Merqury: reference-free quality and phasing assessment for genome assemblies. bioRxiv. doi:10.1101/2020.03.15.992941

Roach, M. J., Schmidt, S. A., & Borneman, A. R. (2018). Purge Haplotigs: allelic contig reassignment for third-gen diploid genome assemblies. BMC Bioinformatics, 19(1), 460. doi:10.1186/s12859-018-2485-7

Robinson, J. A., Brown, C., Kim, B. Y., Lohmueller, K. E., & Wayne, R. K. (2018). Purging of strongly deleterious mutations explains long-term persistence and absence of inbreeding depression in island foxes. Current Biology, 28(21), 3487–3494 e3484. doi:10.1016/j.cub.2018.08.066

Robinson, J. A., Ortega-Del Vecchyo, D., Fan, Z., Kim, B. Y., vonHoldt, B. M., Marsden, C. D., … Wayne, R. K. (2016). Genomic flatlining in the endangered island fox. Current Biology, 26(9), 1183–1189. doi:10.1016/j.cub.2016.02.062

Robinson, J. A., Raikkonen, J., Vucetich, L. M., Vucetich, J. A., Peterson, R. O., Lohmueller, K. E., & Wayne, R. K. (2019). Genomic signatures of extensive inbreeding in Isle Royale wolves, a population on the threshold of extinction. Science Advances, 5(5), eaau0757. doi:10.1126/sciadv.aau0757

Rodriguez-Perez, M. Y., Aurioles-Gamboa, D., Sanchez-Velasco, L., Lavin, M. F., & Newsome, S. D. (2018). Identifying critical habitat of the endangered vaquita (Phocoena sinus) with regional delta C-13 and delta N-15 isoscapes of the Upper Gulf of California, Mexico. Marine Mammal Science, 34(3), 790–805. doi:10.1111/mms.12483

Rohling, E. J., Hibbert, F. D., Williams, F. H., Grant, K. M., Marino, G., Foster, G. L., … Yokoyama, Y. (2017). Differences between the last two glacial maxima and implications for ice-sheet, delta O-18, and sea-level reconstructions. Quaternary Science Reviews, 176, 1–28. doi:10.1016/j.quascirev.2017.09.009

Rojas-Bracho, L., Gulland, F. M. D., Smith, C. R., Taylor, B., Wells, R. S., Thomas, P. O., … Walker, S. (2019). A field effort to capture critically endangered vaquitas *Phocoena sinus* for protection from entanglement in illegal gillnets. Endangered Species Research, 38, 11–27. doi:10.3354/esr00931

Rojas-Bracho, L., & Reeves, R. R. (2013). Vaquitas and gillnets: Mexico’s ultimate cetacean conservation challenge. Endangered Species Research, 21(1), 77–87. doi:10.3354/esr00501

Rojas-Bracho, L., & Taylor, B. L. (1999). Risk factors affecting the vaquita (*Phocoena sinus*). Marine Mammal Science, 15(4), 974–989. doi:10.1111/j.1748-7692.1999.tb00873.x

Rosel, P. E., Haygood, M. G., & Perrin, W. F. (1995). Phylogenetic relationships among the true porpoises (Cetacea:Phocoenidae). Molecular Phylogenetics and Evolution, 4(4), 463–474. doi:10.1006/mpev.1995.1043

Rosel, P. E., & Rojas-Bracho, L. (1999). Mitochondrial DNA variation in the critically endangered vaquita *Phocoena sinus* Norris and Macfarland, 1958. Marine Mammal Science, 15(4), 990–1003. doi:10.1111/j.1748-7692.1999.tb00874.x

Siddall, M., Rohling, E. J., Almogi-Labin, A., Hemleben, C., Meischner, D., Schmelzer, I., & Smeed, D. A. (2003). Sea-level fluctuations during the last glacial cycle. Nature, 423(6942), 853–858. doi:10.1038/nature01690

Smit, A., Hubley, R., & Green, P. (2013-2015). RepeatMasker Open 4.0: http://www.repeatmasker.org.

Springer, M. S., Emerling, C. A., Fugate, N., Patel, R., Starrett, J., Morin, P. A., … Gatesy, J. (2016a). Inactivation of cone-specific phototransduction genes in rod monochromatic cetaceans. Frontiers in Ecology and Evolution, 4, Article No. 61. doi:10.3389/fevo.2016.00061

Springer, M. S., Starrett, J., Morin, P. A., Hayashi, C., & Gatesy, J. (2016b). Inactivation of C4orf26 in toothless placental mammals. Molecular Phylogenetics and Evolution, 95, 3445. doi:10.1016/j.ympev.2015.11.002

Supple, M. A., & Shapiro, B. (2018). Conservation of biodiversity in the genomics era. Genome Biology, 19(131). doi:10.1186/s13059-018-1520-3

Taylor, B. L., Chivers, S. J., Larese, J., & Perrin, W. F. (2007). Generation length and percent mature estimates for IUCN assessments of cetaceans (Administrative Report LJ-07-01). Retrieved from https://swfsc.noaa.gov/publications/TM/SWFSC/NOAA-TM-NMFS-SWFSC-550.pdf

Taylor, B. L., & Rojas-Bracho, L. (1999). Examining the risk of inbreeding depression in a naturally rare cetacean, the vaquita (*Phocoena sinus*). Marine Mammal Science, 15(4), 1004–1028. doi:10.1111/j.1748-7692.1999.tb00875.x

Taylor, B. L., Wells, R. S., Olson, P. A., Brownell, R. L., Gulland, F. M. D., Read, A. J., … Rojas-Bracho, L. (2019). Likely annual calving in the vaquita, *Phocoena sinus:* A new hope? Marine Mammal Science, 35(4), 1603–1612. doi:10.1111/mms.12595

Thomas, L., Jaramillo-Legorreta, A., Cardenas-Hinojosa, G., Nieto-Garcia, E., Rojas-Bracho, L., Ver Hoef, J. M., … Tregenza, N. (2017). Last call: Passive acoustic monitoring shows continued rapid decline of critically endangered vaquita. Journal of the Acoustical Society of America, 142(5), E1512–El517. doi:10.1121/1.5011673

Tollis, M., Robbins, J., Webb, A. E., Kuderna, L. F. K., Caulin, A. F., Garcia, J. D., … Maley, C. C. (2019). Return to the sea, get huge, beat cancer: An analysis of cetacean genomes including an assembly for the humpback whale (*Megaptera novaeangliae*). Molecular Biology and Evolution, 36(8), 1746–1763. doi:10.1093/molbev/msz099

Tunstall, T., Kock, R., Vahala, J., Diekhans, M., Fiddes, I., Armstrong, J., … Steiner, C. C. (2018). Evaluating recovery potential of the northern white rhinoceros from cryopreserved somatic cells. Genome Research, 28(6), 780–788. doi:10.1101/gr.227603.117

Van Cise, A. M., Baird, R. W., Baker, C. S., Cerchio, S., Claridge, D., Fielding, R., … Morin, P. A. (2019). Oceanographic barriers, divergence, and admixture: Phylogeography and taxonomy of two putative subspecies of short-finned pilot whale. Molecular Ecology, 28, 2886–2902. doi:10.1111/mec.15107

Waterhouse, R. M., Seppey, M., Simao, F. A., Manni, M., Ioannidis, P., Klioutchnikov, G., … Zdobnov, E. M. (2017). BUSCO applications from quality assessments to gene prediction and phylogenomics. Molecular Biology and Evolution, 35(3), 543–548. doi:10.1093/molbev/msx319

Westbury, M. V., Hartmann, S., Barlow, A., Wiesel, I., Leo, V., Welch, R., … Hofreiter, M. (2018). Extended and continuous decline in effective population size results in low genomic diversity in the world’s rarest hyena species, the brown hyena. Molecular Biology & Evolution, 35(5), 1225–1237. doi:10.1093/molbev/msy037

Westbury, M. V., Petersen, B., Garde, E., Heide-Jorgensen, M. P., & Lorenzen, E. D. (2019). Narwhal genome reveals long-term low genetic diversity despite current large abundance size. iScience, 15, 592–599. doi:10.1016/j.isci.2019.03.023

Xue, Y., Prado-Martinez, J., Sudmant, P. H., Narasimhan, V., Ayub, Q., Szpak, M., … Scally, A. (2015). Mountain gorilla genomes reveal the impact of long-term population decline and inbreeding. Science, 348(6231), 242–245. doi:10.1126/science.aaa3952

Yim, H.-S., Cho, Y. S., Guang, X., Kang, S. G., Jeong, J.-Y., Cha, S.-S., … Lee, J.-H. (2014). Minke whale genome and aquatic adaptation in cetaceans. Nature Genetics, 46, 88–92. doi:10.1038/ng.2835

Zhou, X., Guang, X., Sun, D., Xu, S., Li, M., Seim, I., … Yang, G. (2018). Population genomics of finless porpoises reveal an incipient cetacean species adapted to freshwater. Nature Communications, 9(1), Article number: 1276. doi:10.1038/s41467-018-03722-x

