## Supplemental materials for "Reference genome and demographic history of the most endangered marine mammal, the vaquita"

### Table S1: BUSCO scores

BUSCO scores of the vaquita reference genome primary haplotype based on the BUSCO v3.1.0 laurasiatheria_odb9 and mammalia_odb9 data sets.

| Category | Laurasiatheria No. | Percentage | Mammalia No. | Percentage |
| --- | --- | --- | --- | --- |
| Complete | 5621 | 89.9% | 3758 | 91.6% |
| Complete and single-copy | 5558 | 88.9% | 3714 | 90.5% |
| Complete and duplicated | 63 | 1.0% | 44 | 1.1% |
| Fragmented | 268 | 4.3% | 188 | 4.6% |
| Missing | 364 | 5.8% | 158 | 3.8% |
| Total BUSCO groups searched | 6253 |  | 4104 |  |

### Table S2. Annotation

Partial annotation summary from NCBI *Phocoena sinus* annotation release 100 (https://www.ncbi.nlm.nih.gov/genome/annotation_euk/Phocoena_sinus/100/) based on the NCBI Eukaryotic Genome Annotation Pipeline (v. 8.3). A) Counts for categories of genes and pseudogenes. B) Counts for categories of CDS's. C) Length summaries for gene categories. D) Count of annotated proteins searched with BLASTP against the UniProtKB/Swiss-Prot curated proteins, with match above query or target coverage threshold (90% and 50%) by species for annotated cetacean species genomes in the NCBI database.

| A) |  |
| --- | --- |
| Genes and pseudogenes | 26,497 |
| protein-coding | 19,069 |
| non-coding | 4,238 |
| transcribed pseudogenes | 21 |
| non-transcribed pseudogenes | 3,131 |
| genes with variants | 10,419 |
| immunoglobulin/T-cell receptor gene segments | 38 |

| B) |  |
| --- | --- |
| CDSs | 52,320 |
| fully-supported | 50,497 |
| with > 5% ab initio | 1,170 |
| partial | 100 |
| with major correction(s) | 2,264 |
| known RefSeq (NP_) | 0 |
| model RefSeq (XP_) | 52,282 |

| C) |  |  |  |  |  |
| --- | --- | --- | --- | --- | --- |
| Feature | Count | Mean length (bp) | Median length (bp) | Min length (bp) | Max length (bp) |
| Genes | 23,307 | 46,677 | 14,250 | 55 | 2,234,275 |
| CDSs | 52,282 | 1,972 | 1,494 | 96 | 103,041 |
| Exons | 249,232 | 348 | 139 | 1 | 23,575 |
| In coding transcripts (NM_/XM_) | 235,846 | 332 | 138 | 1 | 23,575 |
| In non-coding transcripts (NR_/XR_) | 35,561 | 374 | 134 | 2 | 17,001 |
| Introns | 217,934 | 6,182 | 1,448 | 30 | 929,498 |
| In coding transcripts (NM_/XM_) | 208,808 | 6,131 | 1,437 | 30 | 929,498 |
| In non-coding transcripts (NR_/XR_) | 30,917 | 5,498 | 1,473 | 32 | 485,878 |

| D) |  |  | Percent of 19069 coding genes |  |
| --- | --- | --- | --- | --- |
| UniProtKB/Swiss-Prot alignment: | 90% coverage | 50% coverage | 90% coverage | 50% coverage |
| Phocoena Sinus (% of target) | 17081 | 18450 | 89.6% | 96.8% |
| Pocoena sunus (% of query) | 17086 | 18748 | 89.6% | 98.3% |
| Orcinus orca (% of target) | 16237 | 17683 | 85.1% | 92.7% |
| Orcinus orca (% of query) | 16626 | 17890 | 87.2% | 93.8% |
| Tursiops truncatus (% of target) | 11529 | 14801 | 60.5% | 77.6% |
| Tursiops truncatus (% of query) | 13779 | 16701 | 72.3% | 87.6% |
| Neophocaena asiaeorientalis (% of target) | 15149 | 17447 | 79.4% | 91.5% |
| Neophocaena asiaeorientalis (% of query) | 16065 | 17977 | 84.2% | 94.3% |

### Table S3. Genome-wide heterozygosity

Genome-wide heterozygosity values used for Figure 4, from Robinson et al. (2016) unless otherwise stated as “this study”.

| Common Name | Species | IUCN Status | Observed pi | Source | DOI |
| --- | --- | --- | --- | --- | --- |
| Island fox SanNicolas | Urocyon littoralis | NT | 1.42E-05 | Robinson_2016 | 10.1016/j.cub.2016.02.062 |
| Vaquita | Phocoena sinus | CR | 0.00010 | this study |  |
| Baiji | Lipotes vexillifer | CR | 0.00012 | Zhou_2013 | 10.1038/ncomms3708 |
| Brown Hyena | Parahyaena brunnea | NT | 0.00012 | Westbury et al., 2018 | 10.1093/molbev/msy037 |
| Narwhal | Monodon monoceros | LC | 0.00014 | Westbury et al., 2019 | 10.1016/j.isci.2019.03.023 |
| Iberian lynx | Lynx pardinus | CR | 0.00018 | Westbury et al., 2018 | 10.1093/molbev/msy037 |
| Cheetah | Acinonyx jubatus | VU | 0.00020 | Dobrynin_2015 | 10.1186/s13059-015-0837-4 |
| Killer whale | Orcinus orca | NE | 0.00021 | Westbury et al., 2018 | 10.1093/molbev/msy037 |
| Snow leopard | Panthera uncia | VU | 0.00023 | Cho_2013 | 10.1038/ncomms3433 |
| Southern sea otter | Enhydra lutris nereis | EN | 0.00027 | Beichman_2019 | 10.1093/molbev/msz101 |
| Beluga whale | Delphinapterus leucas | LC | 0.00029 | Westbury et al., 2019 | 10.1016/j.isci.2019.03.023 |
| Northern sea otter | Enhydra lutris kenyoni | EN | 0.00031 | Beichman_2019 | 10.1093/molbev/msz101 |
| Tasmanian devil | Sarcophilus harrisii | EN | 0.00032 | Miller_2011 | 10.1073/pnas.1102838108 |
| White African lion | Panthera leo | VU | 0.00048 | Cho_2013 | 10.1038/ncomms3433 |
| Amur tiger (TaeGeuk) | Panthera tigris altaica | EN | 0.00049 | Cho_2013 | 10.1038/ncomms3433 |
| African lion | Panthera leo | VU | 0.00058 | Cho_2013 | 10.1038/ncomms3433 |
| Minke whale | Balaenoptera acutorostrata | LC | 0.00061 | Yim_2014 | 10.1038/ng.2835 |
| Giant otter | Pteronura brasiliensis | EN | 0.00063 | Beichman_2019 | 10.1093/molbev/msz101 |
| Eastern lowland gorilla | Gorilla beringei graueri | CR | 0.00064 | Xue_2015 | 10.1126/science.aaa3952 |
| Orang-Utan (Bornean) | Pongo pygmaeus | CR | 0.00065 | Locke_2011 | 10.1038/nature09687 |
| Mountain gorilla | Gorilla beringei beringei | CR | 0.00065 | Xue_2015 | 10.1126/science.aaa3952 |
| Naked mole rat | Heterocephalus glaber | LC | 0.00068 | Kim_2011 | 10.1038/nature10533 |
| White Bengal tiger | Panthera tigris tigris | EN | 0.00073 | Cho_2013 | 10.1038/ncomms3433 |
| African green monkey | Chlorocebus aethiops aethiops | LC | 0.00080 | Warren_2015 | 10.1101/gr.192922.115 |
| Finless porpoise | Neophocaena phocaenoides | VU | 0.00086 | Yim_2014 | 10.1038/ng.2835 |
| Indo-Pacific finless porpoise | Neophocaena phocaenoides | VU | 0.00093 | this study (based on data from Zhou et al., 2018) | 10.1038/s41467-018-03722-x |
| Chimpanzee W African | Pan troglodytes verus | CR | 0.00095 | The Chimpanzee Sequencing and Analysis Consortium 2005 | 10.1038/nature04072 |
| Cross River gorilla | Gorilla gorilla diehli | CR | 0.00099 | Xue_2015 | 10.1126/science.aaa3952 |
| Human | Homo sapiens | NE | 0.00104 | Corbett-Detig_2015 | 10.1371/journal.pbio.1002112 |
| Yangtze finless porpoise | Neophocaena asiaeorientalis asiaeorientalis | CR | 0.00105 | this study (based on data from Zhou et al., 2018) | 10.1038/s41467-018-03722-x |
| Orang-Utan (Sumatran) | Pongo abelii | CR | 0.00120 | Locke_2011 | 10.1038/nature09687 |
| Gray fox | Urocyon cinereoargenteus | LC | 0.00120 | Robinson_2016 | 10.1016/j.cub.2016.02.062 |
| Cow | Bos taurus | NE | 0.00121 | Corbett-Detig_2015 | 10.1371/journal.pbio.1002112 |
| Narrow-ridged finless porpoise | Neophocaena asiaeorientalis | EN | 0.00134 | this study (based on data from Zhou et al., 2018) | 10.1038/s41467-018-03722-x |
| Giant panda | Ailuropoda melanoleuca | VU | 0.00135 | Leffler_2012 | 10.1371/journal.pbio.1001388 |
| Bottlenose dolphin | Tursiops truncatus | LC | 0.00142 | Yim_2014 | 10.1038/ng.2835 |
| Western lowland gorilla | Gorilla gorilla gorilla | CR | 0.00144 | Xue_2015 | 10.1126/science.aaa3952 |
| Gray wolf | Canis lupus | LC | 0.00149 | Corbett-Detig_2015 | 10.1371/journal.pbio.1002112 |
| Fin whale | Balaenoptera physalus | EN | 0.00151 | Yim_2014 | 10.1038/ng.2835 |
| Chimpanzee C African | Pan troglodytes troglodytes | EN | 0.00176 | The Chimpanzee Sequencing and Analysis Consortium 2005 | 10.1038/nature04072 |
| Vervet monkey | Chlorocebus aethiops pygerythrus | LC | 0.00180 | Warren_2015 | 10.1101/gr.192922.115 |
| Olive baboon | Papio anubis | LC | 0.00189 | Corbett-Detig_2015 | 10.1371/journal.pbio.1002112 |
| Blue whale | Balaenoptera musculus | EN | 0.00210 | this study (based on data from Bukhman et al., in prep.) | Bukhman et al., in prep. |
| Bighorn sheep | Ovis canadensis | LC | 0.00222 | Corbett-Detig_2015 | 10.1371/journal.pbio.1002112 |
| Sperm whale | Physeter macrocephalus | VU | 0.00228 | this study (based on data from Fan et al., 2019) | 10.1111/1755-0998.13003 |
| Rhesus macaque | Macaca mulatta | LC | 0.00287 | Corbett-Detig_2015 | 10.1371/journal.pbio.1002112 |
| Przewalski's horse | Equus ferus przewalskii | EN | 0.00363 | Corbett-Detig_2015 | 10.1371/journal.pbio.1002112 |
| Wild boar | Sus scrofa | LC | 0.00441 | Corbett-Detig_2015 | 10.1371/journal.pbio.1002112 |
| House mouse (SE Asia) | Mus musculus castaneus | LC | 0.00809 | Corbett-Detig_2015 | 10.1371/journal.pbio.1002112 |

### Figure S1. Coalescent model

Distribution of -logL values from A) the initial 20,000 scenarios, and B) the 10,000 scenarios where -logL was less than 6000.

A)

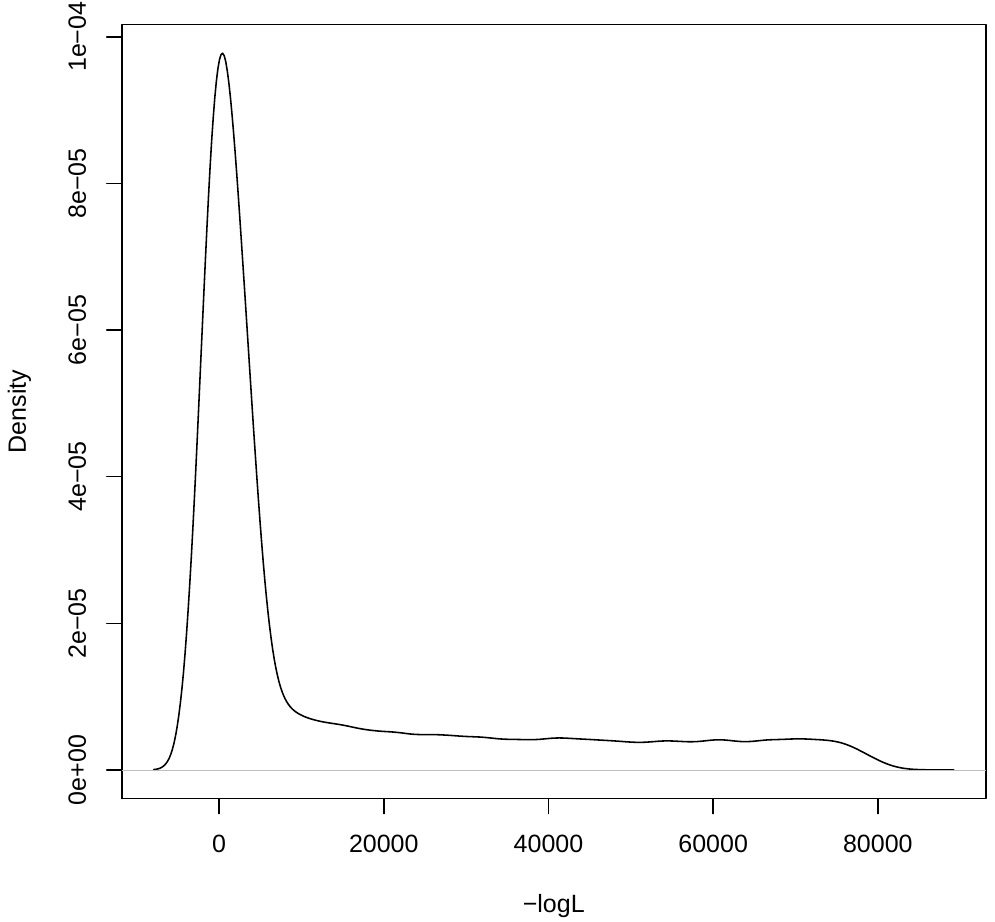

B)

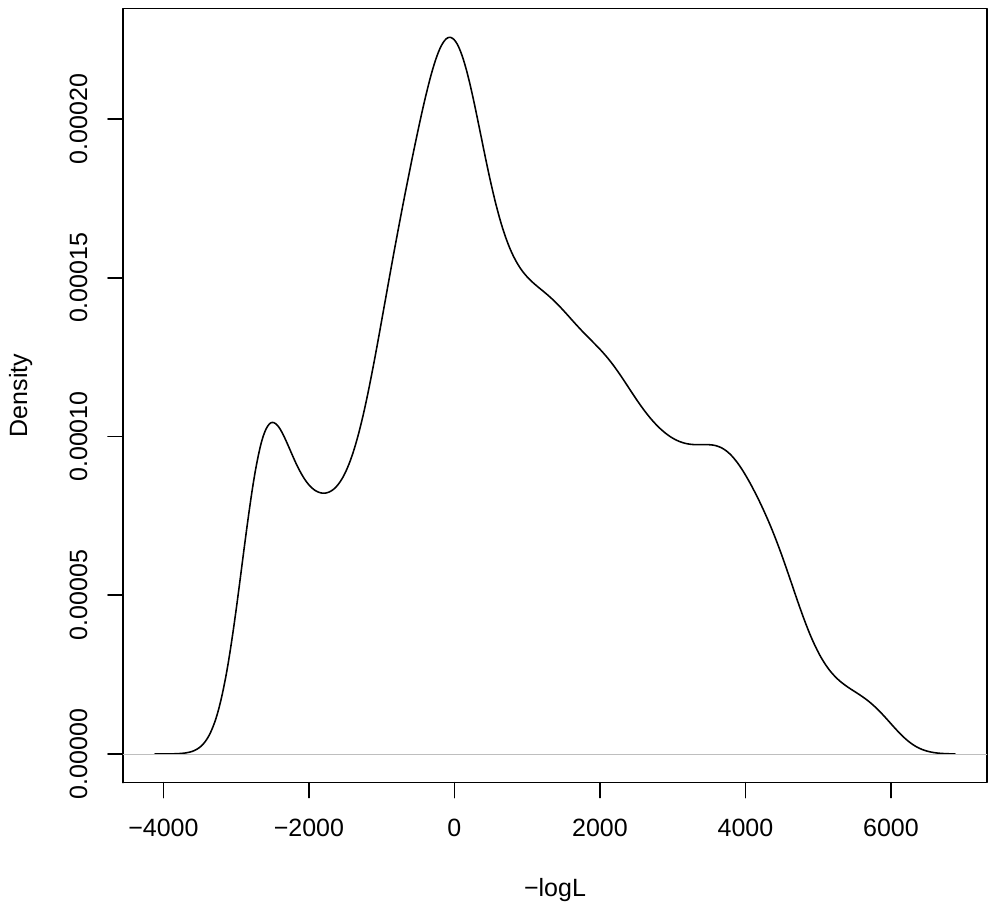
